# A TIR-SAVED effector mediates antiviral immunity via a conserved host signal

**DOI:** 10.1101/2025.06.19.660554

**Authors:** Deepak Kumar Choudhary, Dana Vassover, Himani Singla, Hadar Shukrun, Uri Gophna

## Abstract

Cyclic oligonucleotide-based anti-phage signalling systems (CBASS) are widespread prokaryotic antiviral defense mechanisms that function through coordinated cyclase-effector interactions. Upon sensing viral infection, the cyclase produces a signal molecule that activates effector function and causes cell dormancy or death. However, the evolutionary origins and functional independence of CBASS components remain unclear. Type II CBASS systems commonly employ TIR-SAVED domain effector proteins that deplete cellular NAD^+^ during viral infection. Here, we demonstrate that a TIR-SAVED effector protein can operate as a standalone antiviral defense, causing significant growth inhibition and approximately 50% viral clearance during infection in the complete absence of its cognate cyclase. Remarkably, we show that the TIR-SAVED effector can sense cyclic di-AMP, a conserved second messenger produced by the host diadenylate cyclase DacZ, when the canonical CBASS signal is absent. This antiviral activity was associated with depletion of cellular NAD^+^ and required intact conserved functional residues within both the TIR and SAVED domains. These findings reveal a novel mechanism of antiviral signalling that expands the functional repertoire of CBASS. They also provide insights into the modular evolution of complex prokaryotic immune systems, suggesting that what are now CBASS effectors might have evolved as independent defense components before being integrated into multi-protein systems.

## Introduction

Bacterial and archaeal cells employ diverse antiviral defense mechanisms to combat viral infections. Cyclic-oligonucleotide-based anti-phage signalling systems (CBASS) are of particular interest, since they are common in prokaryotes, and are the ancestors of the animal innate immune factors OAS and C-GAS Sting^1^. These systems are found in almost 14% of bacteria^2,3^ and can activate various cellular responses upon viral infection, triggering cell death or dormancy before the viruses can spread to sister cells ^2,4–8^. To date, four types of CBASS systems have been identified in bacterial and archaeal populations, with approximately 39% being type II CBASS systems^2^. Typically, each type II CBASS gene cluster is composed of genes encoding a cyclase, one or more effectors, and the ancillary proteins Cap2 and Cap3: a ubiquitin-like protein ligase and a deconjugating enzyme, respectively ^2,7,9,10^.

Many Bacterial type II CBASS Systems defend against viral infections through a TIR-SAVED domain effector protein that depletes cellular NAD^+^ levels. Upon viral infection, a cyclic oligonucleotide is generated by the cyclase and sensed by the SAVED domain of the effector protein, which then enables the TIR domain-based NADase activity by oligomerization, which ultimately leads to cell dormancy or death^7,10^. This coordinated mechanism has been extensively characterized in bacterial systems, where mutation of the cyclase active site completely abolishes CBASS activity, reinforcing the view that the proteins of these systems function as interdependent components^10,11^.

However, most if not all complex biological systems have evolved from pre-existing parts via a process that is often referred to as evolutionary tinkering^12,13^. The evolutionary origins and potential functional independence of CBASS components remain unknown. TIR domain proteins are ancient and widely distributed across prokaryotic antiviral systems ^2,7,14–20^, existing both as standalone defense proteins and as components of complex multi-protein systems^14^. Similarly, SAVED domains are found across diverse immunity contexts, including various CBASS types and type III CRISPR systems, suggesting potential functional versatility beyond their canonical signalling roles^2,7,21–25^. This raises the question if the fusion of these domains into one protein preceded their association with the cyclase, and whether CBASS effectors can function independently of their cognate cyclases or recognize alternative signalling molecules beyond their predicted substrates.

If effectors can operate autonomously, this would suggest that complex CBASS systems may have evolved through the recruitment and integration of pre-existing defense components, rather than co-evolution as obligate modules. Furthermore, the ability of effectors to recognize alternative signalling molecules could provide additional layers of antiviral protection and explain the remarkable diversity and widespread distribution of CBASS systems across bacterial and archaeal genomes^2^.

Our previous work demonstrated that an archaeal Type II CBASS system in *Haloferax volcanii* (H-CBASS2) responds to chronic viral infection by inducing growth inhibition rather than triggering cell death^10^, suggesting an alternative mechanism of action in halophilic archaea compared to the typical abortive infection responses observed in bacteria. Here, we investigate the functional independence of TIR-SAVED effector proteins from H-CBASS2. Using the chronically-infecting virus HFPV-1 and the model archaeon *H. volcanii*, we demonstrate that the H-TIR-SAVED effector can function as a standalone antiviral defense system, causing growth inhibition and viral clearance in the complete absence of its cognate cyclase. Remarkably, we show that this independent activity depends on the recognition of cyclic di-AMP, a conserved signal secondary messenger produced by the host diadenylate cyclase DacZ, in the absence of the canonical cyclic trinucleotide signal typically associated with type II CBASS systems that are related in sequence and structure to HCBASS2. Our findings reveal a novel mechanism of antiviral signalling that expands the functional repertoire of known CBASS signals and provides insights into the evolution of complex prokaryotic immune systems.

## Results

### Expression of TIR–SAVED proteins lead to growth inhibition during viral infection, even in the absence of a H-CBASS2 cyclase

We recently showed that the type II CBASS system of *Haloferax* strain Atlit 48N (H-CBASS2) senses chronic virus infection and causes growth inhibition when expressed in *H. volcanii*^10^. This type II CBASS has a TIR-SAVED effector protein, which deplete NAD^+^ following viral infection in the presence of a cyclic nucleotide signal^7,10^.

To investigate the dependence of H-CBASS2 on the signal produced by its cognate cyclase we first cloned H-TIR-SAVED protein (Fig. 1A) separately into the high-copy replicating plasmid pTA927, which contains an inducible promoter, and transformed the constructs into *H. volcanii*, which does not have any CBASS system. We then infected the cells with the HFPV-1, a chronically-infecting virus^26^, and selected virus-infected colonies expressing each individual protein to assess their physiological effects. Interestingly, we observed that virus-infected cells expressing H-TIR-SAVED, despite lacking both the E1-E2-JAB and cyclase proteins, exhibited substantial growth inhibition compared to non-infected cells or those carrying an empty vector (Fig. 1B). A slight growth inhibition was also observed upon H-TIR-SAVED expression, in the absence of virus infection, in *H. volcanii* compared to the control strain (Fig. 1B). We used qPCR to measure viral DNA levels in the supernatant and found that cells expressing H-TIR-SAVED showed a 20-30% reduction in viral DNA compared to control cells (Fig. 1C)

**Figure 1.**
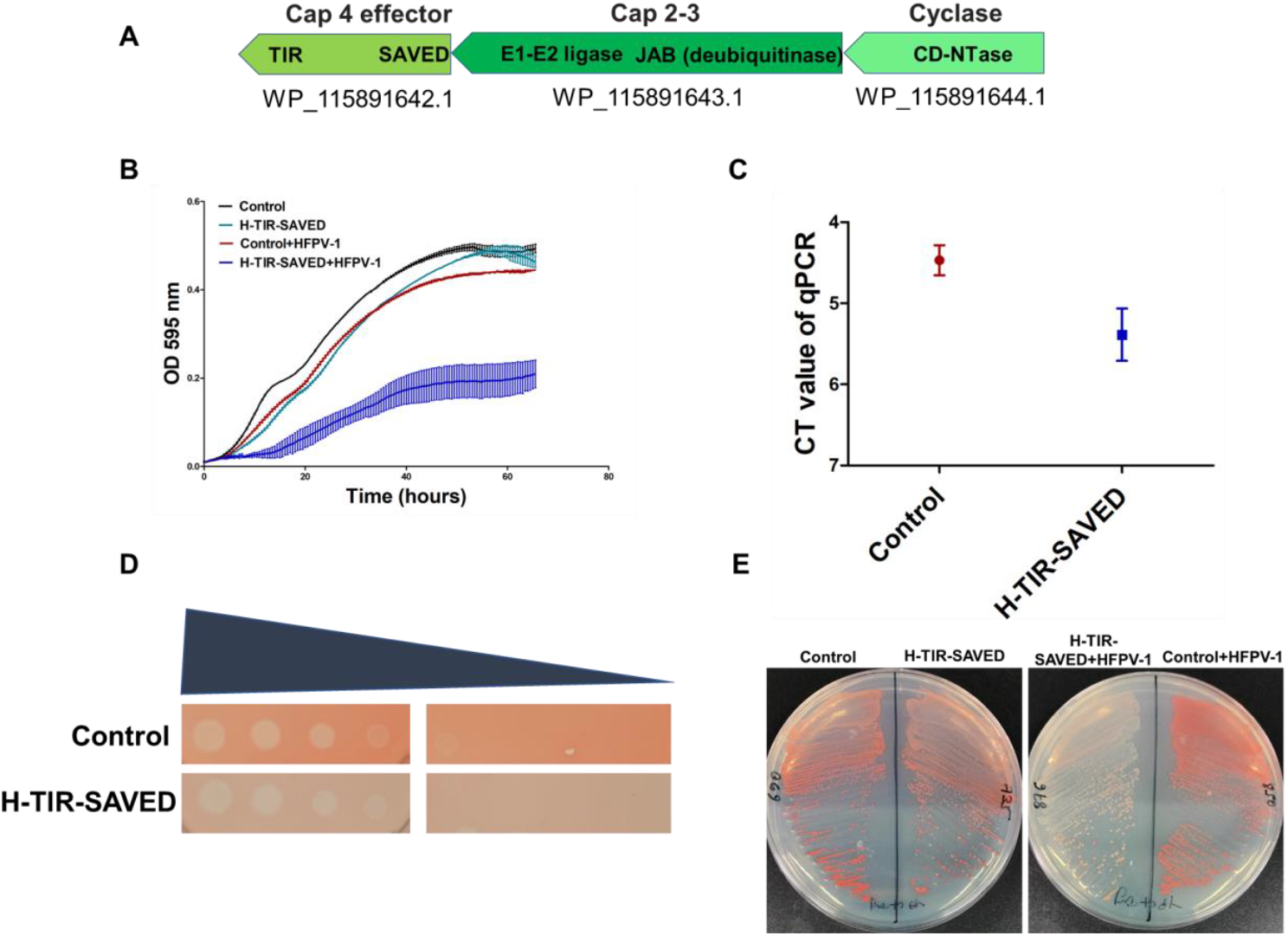
HFPV-1 infection causes growth delay in TIR–SAVED expressing in the absence of the CBASS cyclase. A) The H-CBASS2 operon. The H-CBASS2 TIR-SAVED effector (H-TIR-SAVED) gene was cloned into the pTA927 plasmid, regulated by an inducible tryptophanase promoter, and transformed into *H. volcanii* WR540. HFPV-1 particles were then introduced into the H-TIR-SAVED expressing *H. volcanii* cells, and virus-infected colonies were selected through PCR screening. B) Growth curves of *H. volcanii* cells expressing H-TIR-SAVED under the tryptophan-inducible promoter (cyan) and control cells (empty vector, black), in the absence of viral infection, virus-infected H-TIR-SAVED-expressing cells (blue) under the tryptophan-inducible promoter and infected control cells, with an empty pTA927 (red). Values represent the mean ± SE of at least 3 biologicals replicates, each with three technical replicates. C) Viral DNA concentration in the supernatant of HFPV-1 infected *H. volcanii* cells after 72 hours of growth represented as Ct values determined using primers specific to the HFPV-1 genome (mean ± SE of three biological replicates). D) Plating efficiency of HFPV-1 measured by a plaque assay conducted using 10-fold serial dilutions of virus stock (the left-most dilution is a 10^−4^ virus suspension). H-TIR-SAVED expressing cells, as well as control cells with an empty vector, were mixed with 0.02% top agar and plated onto rich medium. Then, 3 µL of serial dilutions of HFPV-1 were spotted onto the plates to measure plaquing efficiency. The images shown are representative of at least three biological replicates. E) Bleaching of virus-infected H-TIR-SAVED-expressing *H. volcanii* colonies. Representative images of colony color for *H. volcanii* cells infected with HFPV-1 using *H. volcanii* strains expressing H-TIR-SAVED, either with or without viral infection, along with a control containing an empty vector. A 20 µL log-phase culture was streaked onto fresh plates, which were incubated at 45°C for 4-5 days. After growth became visible, the plates were stored at room temperature for 3-4 weeks, and images were captured.

Next, we performed live-dead staining assays on logarithmic cultures of HFPV-1-infected cells expressing H-TIR-SAVED proteins. In agreement with our previous findings on the full CBASS system^10^, no dead cells were observed in these cultures (Extended Fig. 1a). As expected, H-TIR-SAVED expression also did not affect the plaquing efficiency of HFPV-1(Fig. 1D). Similar to the full-length type II CBASS system^10^, the expression of H-TIR-SAVED caused a growth delay but did not induce cell death, reinforcing the idea that type II CBASS systems and their components do not function as a typical abortive infection mechanism, at least not during chronic HFPV-1 infection. We then investigated the long-term effects of viral infection by exposing cells expressing H-TIR-SAVED to room temperature conditions for 3-4 weeks. The virus-infected colonies of cells expressing the H-TIR-SAVED (NAD-depleting) effector exhibited noticeable bleaching compared to infected H-TIR-SAVED-negative control cells (Fig. 1E), similar to the results obtained previously with the full system^10^. As expected, subsequent live-dead assays after long-term virus exposure revealed a higher proportion of dead cells in virus-infected H-TIR-SAVED culture compared to infected controls (Extended Fig. 1b). We also observed that HFPV-1 infection caused more severe growth delay in cells expressing H-TIR-SAVED compared to those expressing full-length H-CBASS2^10^ (Extended Fig. 2a). Additionally, virus-infected H-TIR-SAVED cells bleached more rapidly than cells expressing full-length H-CBASS2 (Extended Fig. 2b). Taken together, these results confirm that the TIR-SAVED effector is responsible for the bleaching activity previously observed for H-CBASS2, which probably results from depriving cells of NADPH required for the synthesis of isoprenoids, including carotenoids^27–29^.

### The TIR-SAVED effector protein depletes NAD^+^ in virus-infected cells, In the absence of the CBASS cyclase

Previous studies have demonstrated that type II CBASS systems detect viral infection through their cyclase proteins, which then produce cyclic signalling molecules that are sensed by effectors, ultimately triggering cell death or dormancy^7^.Accordingly, our previous results showed that inactivating the cyclase by mutation abolished the H-CBASS2 activity during viral infection^10^. This raised the question of whether the observed growth delay in H-TIR-SAVED-expressing cells was caused by NAD^+^ depletion. To determine whether the H-TIR-SAVED protein was able to deplete cellular NAD^+^ levels in the absence of a cyclase we measured total NAD levels in logarithmic-phase cultures of HFPV-1-infected cells expressing H-TIR-SAVED. Interestingly, we found that total NAD levels decreased in virus-infected H-TIR-SAVED-expressing cells compared to infected controls containing an empty vector (Fig. 2A). In contrast, total NAD levels in infected cells expressing the full H-CBASS2 remained unchanged during logarithmic growth^10^. Moreover, we also observed a substantial reduction in total NAD levels, in infected bleached colonies expressing H-TIR-SAVED in the absence of a virus (Fig. 2B). Notably, when measuring NAD^+^ levels, we observed that infected H-TIR-SAVED-expressing cells showed a two-fold greater decrease compared to cells expressing the full-length H-CBASS2 (Extended Fig. 2c). Next, we generated the H-TIR-SAVED and *Microbacterium ketosireducens* TIR-SAVED structures using AlphaFold 3^30^. We then compared the H-TIR-SAVED structure with the *M. ketosireducens* TIR-SAVED structure using Chimera and created a homology model ^31^, which revealed a high degree of structural similarity (RMSD between 196 pruned atom pairs is 1.188 angstroms; (across all 383 pairs: 3.757) (Fig. 2C). Further analysis identified E80 as the active-site residue in the TIR domain, which we mutated to glutamine (E80Q). Notably, previous studies reported that this mutation in *M. ketosireducens* TIR-SAVED abolished both NAD depletion and phage defense^7,32^. Consistent with these findings, the E80Q mutation in H-TIR-SAVED fully restored NAD levels and partially rescued the growth phenotype (Fig. 2D&E) in H-TIR-SAVED -expressing infected cells. Furthermore, we did not observe bleached colonies in virus-infected cells after introducing this mutation (Extended Fig. 3).

**Figure 2.**
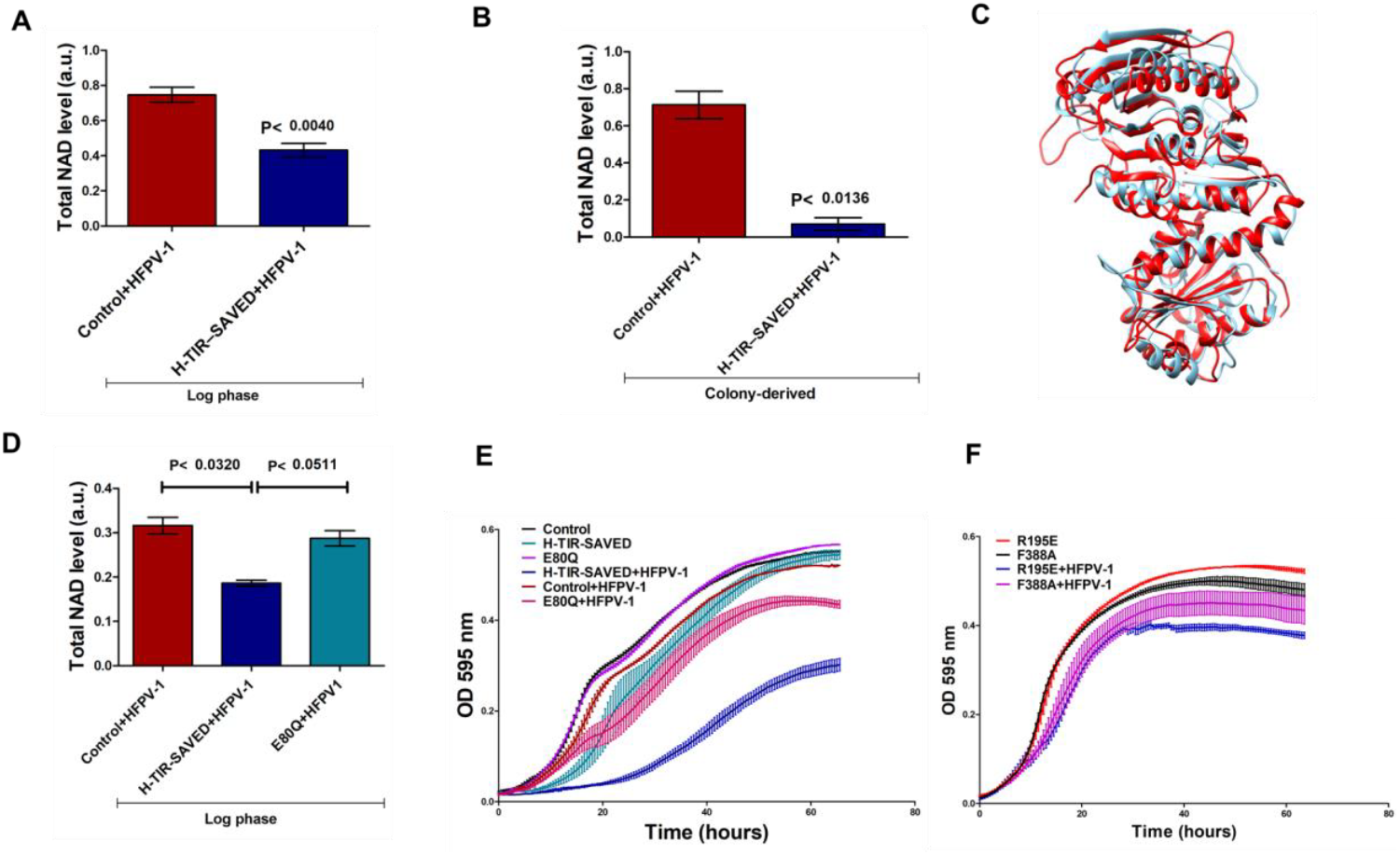
TIR-SAVED protein reduces NAD^+^ levels in virus-infected cells independently of CBASS cyclase activity. A) NAD^+^ levels in *H. volcanii* strains expressing H-TIR-SAVED following virus infection. Log-phase cultures with equal cell numbers were collected, and total NAD^+^ levels were measured using the NAD/NADH Quantification Kit. B) NAD^+^ levels in cells from bleached colonies. Bleached cells incubated for 3–4 weeks were resuspended in fresh medium, and equal protein-normalised biomass from each strain was used to quantify total NAD^+^. A and B: Data represent results from three independent experiments. NAD^+^ values of the different strains were compared using a paired sample t-test. C) 3D structural alignment comparing predicted H-TIR-SAVED and *M. ketosireducens* TIR-SAVED structures. Both structures were generated using AlphaFold 3 and aligned using ChimeraX software (RMSD = 3.757 Å). NAD^+^ levels in *H. volcanii* strains expressing the Q80E mutation in TIR-SAVED protein. Strains were grown to log phase and NAD^+^ levels were measured following the protocol described. Data are representative of three independent experiments. Statistical analysis was performed using a paired t-test. E) Growth curves showing the effect of the E80Q mutation in the H-TIR-SAVED protein. E80Q without infection is shown in purple, and with infection in pink. Control strains are shown in black (uninfected) and red (infected). Wild-type H-TIR-SAVED is shown in cyan (uninfected) and blue (infected). E) Growth curves of H-TIR-SAVED mutants R195E and F388A with and without viral infection. R195E is shown in red (uninfected) and blue (infected), while F388A is shown in black (uninfected) and pink (infected). E and F: Values represent the mean ± SE of at least three biological replicates, each with three technical replicates.

Recent studies have shown that TIR-SAVED oligomerizes and becomes active in the presence of cA3 signalling molecules. This multimerization is crucial for its function, as mutations in the SAVED domain active site disrupt signal binding (specifically to cA3)^7^, rendering the protein monomeric and inactive. We investigated whether structurally analogous amino acids in H-TIR-SAVED would abolish its activity, which would indicate that, like a typical CBASS system, the H-SAVED domain binds specific signalling molecules in response to infection. To test this hypothesis, we introduced R195E and F388A mutations separately into the H-TIR-SAVED active site and found that the F388A mutation partially reversed the growth delay, partially restored NAD levels, and prevented the colony bleaching caused by HFPV-1 infection (Fig. 2F, Extended Fig. 4). This result indicates that, like a typical CBASS system, the SAVED domain must bind a specific signalling molecule in order to be active during infection.

### H-TIR-SAVED proteins sense cyclic di-AMP, a signaling molecule produced by a conserved host enzyme

Since introducing mutations in the signal binding site of H-TIR-SAVED abolished activity, this implied that a signalling molecule is required, which must be generated by a host enzyme in the absence of a cyclase during viral infection. Since H-TIR-SAVED is induced and may adopt an open conformation, it could potentially bind to molecules resembling its predicted substrate cAAG that are somehow produced as a result of membrane stress triggered by viral infection. The obvious candidate in the case of *H. volcanii* is the essential secondary messenger molecule cyclic-di-AMP (c-di-AMP)^33,34^. This molecule is produced by the diadenylate cyclase DacZ and has been shown to be essential, required for osmoregulation and, strikingly, toxic when its concentration rises above a certain threshold within cells^33,34^. Some diadenylate cyclases are found in proximity to CBASS gene clusters, and have been suggested to sense phage infection by responding to membrane stress^35,36^.To test this hypothesis, we used *H. volcanii* strains with reduced and constitutive DacZ expression, exhibiting approximately 50% reduction in activity relative to basal levels^33^. We then introduced the H-TIR-SAVED protein along with HFPV-1 virus into these low c-di-AMP cells. Notably, cells expressing H-TIR-SAVED in these strains showed substantially milder growth inhibition when infected with HFPV-1 (Fig. 3A). Furthermore, we added exogenous c-di-AMP to cells expressing either the full H-CBASS2 and H-TIR-SAVED proteins without viral infection and performed growth analyses. Interestingly, we found that in the presence of external c-di-AMP, the H-TIR-SAVED protein caused growth inhibition even in the absence of virus. We also observed a small effect on the full-length CBASS system (Fig. 3B). Notably, we did not observe any effect of c-di-AMP when we used strains with mutations in the TIR and SAVED domain proteins (Fig. 3C), as expected.

**Figure 3.**
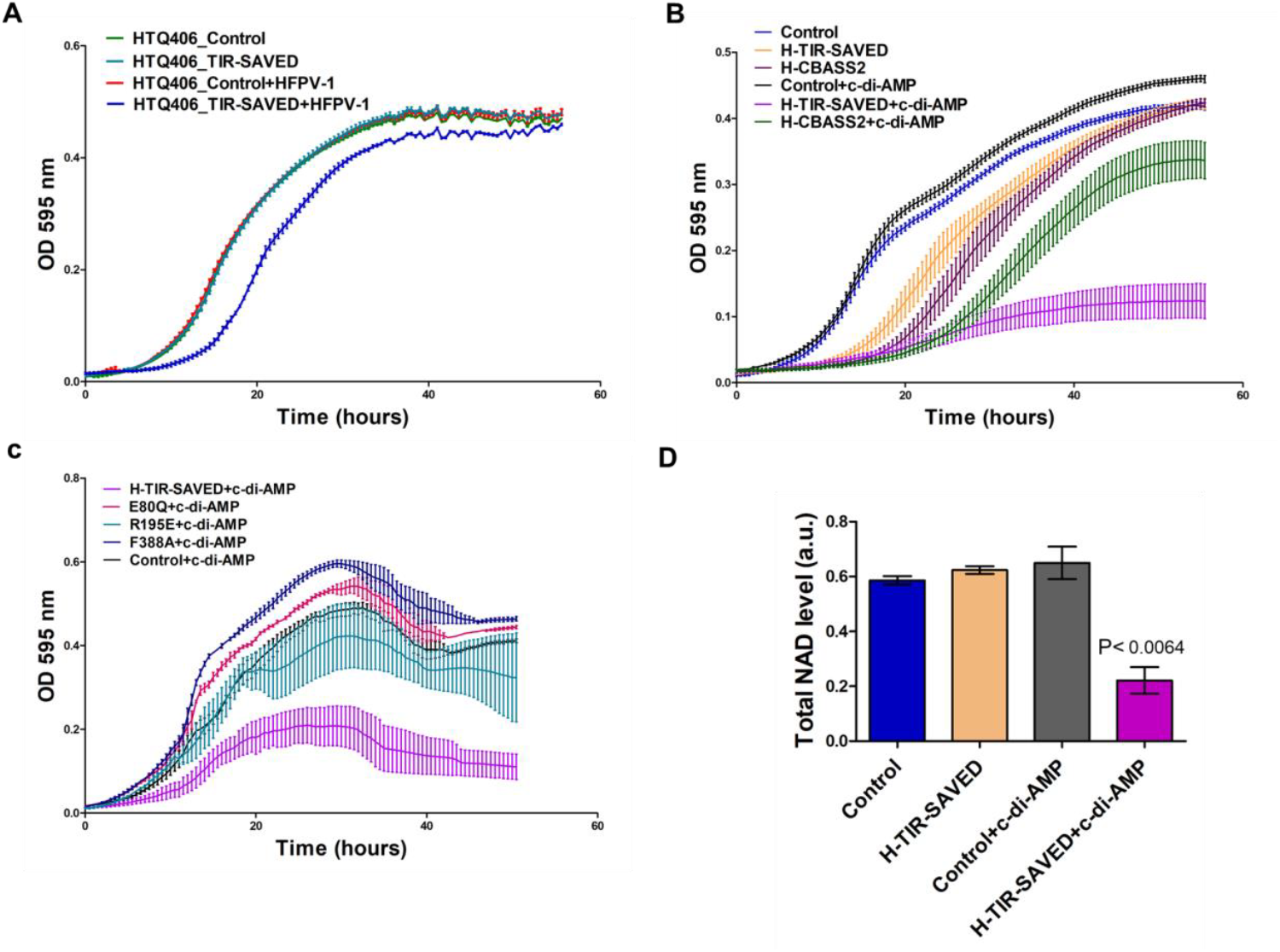
H-TIR-SAVED senses the physiological stress molecule cyclic di-AMP. A) Growth curves of cells of *H. volcanii* strain HTQ406 that has reduced intracellular c-di-AMP levels^33^. H-TIR-SAVED was expressed in HTQ406 cells that were infected with HFPV-1. Infected H-TIR-SAVED-expressing cells are shown in blue, infected controls with empty vector in red, uninfected H-TIR-SAVED-expressing cells in cyan, and uninfected controls with empty vector in green. B) Growth analysis of *H. volcanii* strains expressing H-CBASS2 (green) or H-TIR-SAVED (purple) in the presence of exogenous c-di-AMP, compared to untreated H-CBASS2 or H-TIR-SAVED-expressing cells (dark purple) or yellow, respectively). c-di-AMP-Treated and untreated control cells are shown in black and blue. C) Growth curves of strains carrying mutations in the TIR and SAVED domains in the presence of c-di-AMP. The wild-type H-TIR-SAVED strain with c-di-AMP is shown in purple, and its corresponding control in black. Mutants E80Q (TIR domain), R195E, and F388A (SAVED domain) are shown in pink, cyan, and blue, respectively. Values represent the mean ± SE of at least three biological replicates, each with three technical replicates. D) NAD^+^ levels in *H. volcanii* expressing H-TIR-SAVED following c-di-AMP treatment. Log-phase cultures of c-di-AMP-treated and untreated cells, normalized to equal cell numbers by OD, were collected, and total NAD^+^ levels were measured as described in the Methods. D: Data represent results from at least three independent experiments. NAD^+^ values of the different strains were compared using a paired sample t-test.

Next, we measured total NAD levels in cells expressing H-TIR-SAVED following supplementation with exogenous c-di-AMP. Interestingly, total NAD levels were reduced in H-TIR-SAVED-expressing cells compared to treated control cells containing an empty vector (Fig. 3D), upon c-di-AMP supplementation. Taken together, these results indicate that the H-CBASS TIR-SAVED effectors responds to c-di-AMP signalling, in the absence of its cognate CBASS signal.

### HFPV-1 infection increases cyclic di-AMP levels in *H. volcanii* cells

Having established that H-TIR-SAVED proteins sense c-di-AMP produced by DacZ upon virus infection, we next tested whether virus infection can increase the production of cellular c-di-AMP. We therefore measured c-di-AMP levels in log-phase *H. volcanii* cells with or without HFPV-1 infection. Interestingly, we found that virus-infected cells had almost 25% higher c-di-AMP levels compared to control cells (Fig. 4A). These results demonstrate that virus infection directly stimulates c-di-AMP production in *H. volcanii*, thereby providing the necessary signal for H-TIR-SAVED activation.

**Figure 4.**
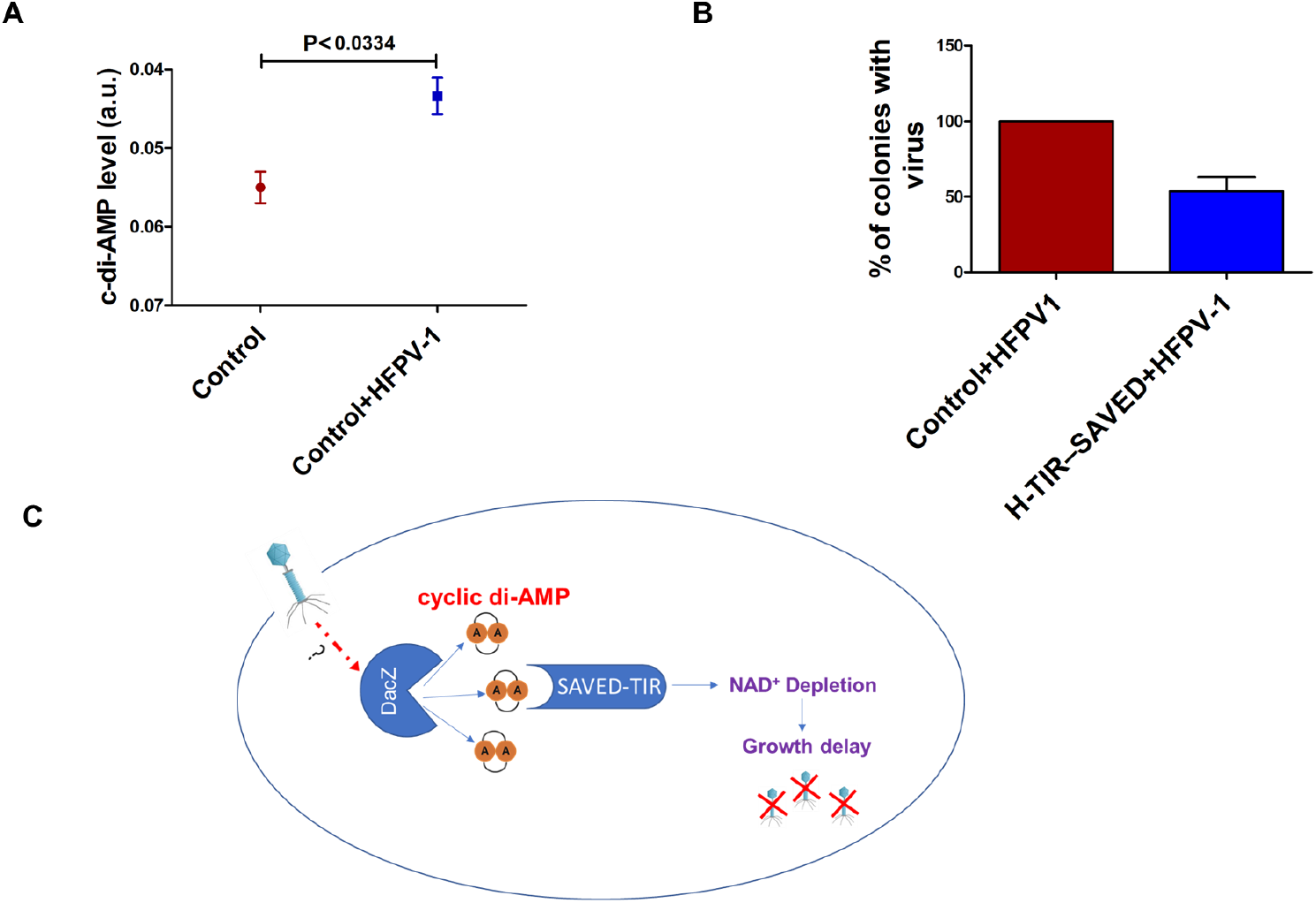
Expression of H-TIR-SAVED leads to clearance of chronic virus infection in *H. volcanii*. **A)** Elevated c-di-AMP Levels in *H. volcanii* following Viral Infection. Log-phase cultures with equal cell numbers were collected, and total c-di-AMP levels were measured using the c-di-AMP ELISA Kit. The absorbance at 450 nm, measured spectrophotometrically, reflects the amount of cyclic di-AMP-HRP tracer bound to the wells and is inversely proportional to the concentration of free c-di-AMP present during the incubation period. B) Ratio of virus-free colonies vs. virus-infected after three passages in rich medium containing tryptophan. *H. volcanii* strain infected with HFPV-1 and expressing H-TIR-SAVED was compared to a virus-infected control strain, and virus presence was determined using colony PCR. C) Proposed model for the mechanism of action of the TIR-SAVED effector in the absence of a cognate cyclase. Upon HFPV-1 infection, the DacZ protein senses membrane stress caused by the presence of the virus and produces c-di-AMP signalling molecules. The SAVED domain detects the c-di-AMP signal, triggering activation of the TIR domain. The activated TIR domain then depletes cellular NAD^+^ levels, leading to growth delay and eventual bleaching and death.

### H-TIR-SAVED clears a chronic virus infection in *H. volcanii* cells

We have observed that expression of full-length CBASS2 (H-CBASS2) is able to clear chronic virus infection after several passages in rich medium^10^. Additionally, expression of H-TIR-SAVED shows a stronger growth inhibition than the H-CBASS2 system (Extended Fig. 2a). Therefore, we next investigated whether H-TIR-SAVED eliminates the virus as effectively as the complete H-CBASS2 system does. We therefore cultured HFPV-1-infected *H. volcanii* cells in liquid medium until late log or stationary phase, comparing cultures either expressing or not expressing H-TIR-SAVED. These cultures were then used to inoculate fresh medium, and this process was repeated for 6 passages. Samples were collected at passages 3 for analysis. We plated diluted cultures on solid media and screened 25-30 colonies from each replicate by PCR to detect the presence of HFPV-1 DNA. Interestingly, virus-infected cells expressing the H-TIR-SAVED protein demonstrated significant antiviral efficacy despite the absence of both E1-E2-JAB and cyclase proteins. By passage 3, these cells showed approximately 50% virus clearance (Fig. 4B). This antiviral effect became even more pronounced by passage 6, where nearly all colonies from H-TIR-SAVED-expressing cultures had undetectable levels of viral DNA across all three biological replicates. In contrast, almost all control colonies that did not express H-TIR-SAVED maintained detectable viral DNA presence (Fig. 4B). Taken together, the independent antiviral and NADase activities, and the effect on growth strongly suggest that the H-TIR-SAVED protein can function as an **independent** immune effector in *H. volcanii* and can respond to viral infection by sensing c-di-AMP signals produced by different cyclase such as DacZ within the cell, rather than being limited to detecting their putative cAAG signal (Fig. 4C).

## Discussion

Here we show that the TIR-SAVED effector of a type II CBASS can act independently as an antiviral defense, and is capable of inducing growth inhibition during infection in the absence of cyclase protein (Fig. 1B). Such independent activity may provide a more robust defense system that is less sensitive to anti-defense mechanisms, such as anti-CBASS (Acb) proteins that sequester signal molecules and inhibit CBASS-based immunity^37,38^. This is consistent with a system that has evolved to have co-dependency between its components. Taken together, these results help explain how complex CBASS systems evolved from their individual components, prior to becoming inter-dependent. TIR-domain proteins are found in many different anti-phage systems, and there are even single protein TIR-domain systems^14^, and thus could have been recruited for defensive roles before becoming associated with a cognate cyclase. Consistent with previous studies in other organisms, we show that the TIR-SAVED effector can cause growth inhibition by depleting NAD^+^ levels in infected cells^7,10^. Interestingly, we found that the expression of TIR-SAVED without the other CBASS components also led to a significant reduction in NAD^+^ levels in virus-infected *H. volcanii* cells (Fig. 2A&B), confirming this domain’s critical role in the observed growth inhibition. The E80Q mutation in the catalytic site of the TIR domain partially reversed the growth phenotype and fully prevented the NAD^+^ depletion caused by virus infection in cells expressing the stand-alone TIR-SAVED protein (Fig. 2D&E).

We further show that the SAVED domain’s ability to mediate antiviral responses depends on specific conserved residues required for signal sensing. The rescue of the growth phenotype and suppression of bleaching by the F388A mutation (which is important for cA3-binding in b bacteria^7^ (Fig. 2F) support a role for the SAVED domain as a signal sensor, likely detecting a cyclic nucleotide signal even in the absence of the canonical cyclase. This is further supported by our observation that H-TIR–SAVED can sense cyclic di-AMP (c-di-AMP), representing a novel twist in the study of CBASS signalling specificity. The substantial rescue of growth inhibition in strains with reduced DacZ expression (Fig. 3A), along with the induction of growth arrest by exogenous c-di-AMP in the absence of viral infection (Fig. 3B&C), provides strong evidence that this effector protein has evolved to recognize endogenous signalling molecules beyond the canonical cA3G produced by CBASS cyclases. These results suggest that while the SAVED domain retains its ligand-binding role, it has broad enough specificity to recognize structurally related cyclic nucleotides beyond its cognate trinucleotide, which is predicted by structural similarity to be cAAG. This expanded signal-recognition capacity may explain the widespread presence of SAVED domains across diverse antiviral immune systems, including various CBASS types as well as type III CRISPR systems^2,5,7,39^.

Our observation that viral infection can increase c-di-AMP levels in *H volcanii* is not really surprising (Fig. 4A). C-di-AMP plays several key roles in bacterial physiology, including responses to osmotic stress and protecting DNA integrity^40–42^. While membrane stress is more typical to early stages of viral infection, at least in bacteria, DNA damage is generally not considered to be phage associated. However, replication of phage genomes often includes DNA open ends, which are indistinguishable from DNA breaks at the molecular level, and could therefore indicate viral replication and trigger a defensive function. Notably, many genes encoding DacZ enzymes naturally co-occur with CBASS systems^35,36^, suggesting a potential link between viral infection, c-di-AMP signalling, and CBASS activation. This also could explain why prokaryotes typically encode a complete CBASS system rather than a standalone TIR-SAVED effector. If TIR-SAVED operated independently based on c-di-AMP signalling alone, it could respond to various physiological stresses in the absence of infection. Such response would not be beneficial for the cells since a response that delays growth and eventually can result in cell death should be reserved exclusively to stopping the spread of viral infection rather triggered during general stress situations.

These findings of cyclase-independent effector function are in contrast to our previous observations that when the whole H-CBASS2 cluster was expressed together, and cyclase activity was abolished by point mutation^10,43^,this resulted in complete abolishment of CBASS activity. This indicates that either the E1-E2 protein, the H-CBASS2 cyclase or both can modulate the effector function until the cyclase produces enough specific signal. The molecular basis for this molecular remains to be elucidated, but one may speculate that the complete system is better for cells than the lone TIR-SAVED effector because it is less likely to respond to spurious membrane stress in a way which is harmful, in the absence of viral infection.

Altogether, our findings suggest that H-TIR-SAVED proteins can function as standalone immune effectors that sense c-di-AMP signals produced by different cellular cyclases to detect viral infection and mediate defense responses. This expands the functional repertoire of CBASS effectors beyond their originally defined signalling context and highlights a more modular, signal-flexible architecture in immunity, which could extend beyond archaea to many Gram-positive bacteria whose genomes encode both *dacZ* and CBASS.

## Methods

### Culture conditions

The Haloferax wild-type strains were regularly cultured at 45°C in either Hv-YPC or Hv-Ca/Hv-Enhanced Ca medium. *Haloferax volcanii* transformants were selected and grown in either Hv-Ca or Hv-Enhanced Ca (Hv-ECa) medium. Thymidine (40 μg/ml) and tryptophan (50 μg/ml) were added when necessary. Bacterial strains were cultured at 37°C in LB medium, or in LB medium supplemented with ampicillin for strains containing plasmids.

### Cloning and mutagenesis

The *Haloferax* 48N TIR-SAVED proteins of the CBASS type II system were cloned into the high-copy replicating plasmid pTA927 (which contains the tryptophanase inducible promoter) using the Gibson assembly method. DNA fragments for the inserts along with the plasmid vectors, were first PCR-amplified using specific primers and either Phusion or KAPA-HiFi DNA polymerase. The PCR-amplified plasmids and DNA fragments were purified using the Wizard® SV Gel and PCR Clean-Up Kit (Promega), followed by DpnI digestion of the purified plasmids. The purified DNA fragments and plasmids were then ligated using the Gibson assembly protocol (Gibson et al., 2009). The constructed plasmids were introduced into *E. coli* DH12S cells through electroporation. The clones were verified using PCR, and the plasmids were subsequently isolated using Sigma-Aldrich’s GenElute™ Plasmid Miniprep Kit. The purified plasmids were then introduced into *H. volcanii* strains following previously published protocols (Halohandbook).

### Mutagenesis

The TIR-dead variant of H-TIR-SAVED cloned into pTA927, was generated by replacing the conserved glutamic acid at position 80 with glutamine using site-directed mutagenesis. Similarly, mutations in the SAVED domain of H-TIR-SAVED, cloned into pTA927, were introduced by substituting the conserved arginine at position 195 with glutamic acid, and the phenylalanine at position 388 with alanine, using site-directed mutagenesis. All the mutation was confirmed through sequencing.

### Transformation with HFPV-1

*H. volcanii* cultures were grown overnight until reaching log phase and collected by centrifugation (6,500 rpm, 5 min). The cell pellet was carefully resuspended in 200 µl of spheroplasting solution (1 M NaCl, 27 mM KCl, 50 mM Tris-HCl, 15% sucrose). The cells were treated with 0.5 M EDTA (pH 8) to remove divalent cations. After 10 minutes at room temperature, concentrated HFPV-1 particles (5-10 µl) were added and incubated for 5 minutes. The suspension was then mixed gently with 250 μL of 60% PEG600 and incubated for 1 hour at room temperature. The cells were washed with 1 ml regeneration solution (Hv-YPC+ media containing 15% sucrose) and resuspended in the same media. The culture was first incubated statically at 28°C for 3 hours, followed by another 3-hour incubation with shaking. The cells were then serially diluted, plated, and individual colonies were screened for successful HFPV-1 infection using virus-specific PCR primers.

### Growth curves

The strains were cultured overnight in Hv-ECa medium supplemented with thymidine and tryptophan until reaching log phase. The cultures were then diluted in fresh Hv-ECa medium containing thymidine and tryptophan to achieve an initial OD of 0.03-0.05. For induction experiments, L-tryptophan was added to a final concentration of 2 mM. The diluted cultures were grown in 96-well plates at 42°C with continuous shaking for 72 hours. Growth was monitored by measuring optical density (OD595nm) at 30-minute intervals using a Biotek ELX808IU-PC microplate reader. Each strain was tested in at least three biological replicates, with each biological replicate consisting of three technical replicates.

### Growth curve with external cyclic di-AMP

Growth experiments were performed using uninfected cells expressing H-TIR-SAVED and a control strain containing the empty plasmid. Each strain was grown overnight in Hv-ECa medium supplemented with thymidine and tryptophan to log phase, then diluted to an OD_600_ of 0.03-0.05 in fresh medium containing 1 µM cyclic di-AMP. The cultures were incubated at 45°C with continuous shaking for 72 hours. Growth was monitored by measuring optical density as described above.

### Growth curve with lower cyclic di-AMP

To examine the effect of cyclic di-AMP, the H-TIR-SAVED gene was first cloned into pTA927. Both this construct and the empty plasmid (control) were transformed into *H. volcanii* strain HTQ406, which has approximately 50% lower intracellular c-di-AMP levels^33^. The HTQ406 strain expressing H-TIR-SAVED was then infected with HFPV-1 virus as described above. The cultures were incubated at 45°C with continuous shaking for 72 hours. Growth was monitored by measuring optical density as described above.

### Live/Dead staining

Cell viability in virus-infected *H. volcanii* strains expressing H-TIR-SAVED was assessed using the LIVE/DEAD bacterial Viability Kit (Thermo-Fisher, Ref-L13152). Overnight cultures were diluted in fresh medium to an OD of 0.03-0.05 and grown at 45°C with shaking until reaching logarithmic phase. Aliquots of 100 µl from each culture were transferred to 1 ml Eppendorf tubes. The cells were stained by adding 10 μl each of Propidium iodide and SYTO9 dyes from their respective stock solutions. After incubating in darkness for 15-20 minutes to allow dye uptake, 10 μl of each stained culture was examined under a Leica sp8 confocal microscope at 20x and 40x magnification. For analysing dead cells in bleached colonies, cells were collected directly from plates containing bleached colonies and suspended in 100 μl of Hv-ECa medium. The cell suspension was washed twice with Hv-ECa medium before proceeding with the previously described microscopy staining and imaging protocol.

### Plate colony pigmentation assays

Strains were grown overnight until reaching log phase, then diluted in fresh medium to OD 0.05. From each diluted culture, 20 µl was streaked onto fresh plates and incubated at 45°C for 4-5 days. After initial growth, the plates were stored at room temperature for 3-5 weeks, with images documented every three days.

### NADase assay

Total cellular NAD was quantified using the NAD/NADH Quantification Kit (Sigma-Aldrich, MAK037). Cultures were grown to late log phase, and equivalent cell numbers (normalized by optical density) were collected and washed twice with cold Hv-Ca media. The cells were pelleted by centrifugation (3000 rpm, 5 min) in 1.5 ml microcentrifuge tubes. Each pellet was resuspended in 400 µl of NADH/NAD extraction buffer and lysed using two rounds of sonication. The lysates were centrifuged (13,000 rpm, 10 min), and the supernatants were filtered through Amicon 10 kDa cut-off spin filters to remove proteins. For the assay, 50 µl of each deproteinized sample was transferred to a 96-well plate. A master reaction mixture (98 µl NAD cycling buffer + 2 µl NAD cycling enzyme mix) was prepared, and 100 µl was added to each well. After 5 minutes of incubation at room temperature, 10 µl of NADH Developer was added per well. The plates were incubated at room temperature for 1-2 hours, followed by absorbance measurement at 450 nm. Bleached colonies were collected directly from plates and suspended in 1 ml of cold Hv-Ca media, followed by two washing steps. The cells were pelleted by centrifugation (3000 rpm, 5 min) in 1.5 ml microcentrifuge tubes. Each pellet was resuspended in 400 µl of NADH/NAD extraction buffer and lysed using two rounds of sonication. Protein concentration in the lysates was determined using the Bradford assay before proceeding with the NAD quantification protocol as previously described.

### Cyclic di-AMP level quantification

Total cellular c-di-AMP levels were quantified using the c-di-AMP Quantification Kit (Cayman chemical, Cyclic di-AMP ELISA Kit Item No. 501960). Cultures were grown to log phase, and equivalent cell numbers (normalized by optical density) were collected and washed with cold Hv-Ca media. The cells were pelleted by centrifugation (5,000 rpm, 5 min) in 1.5 ml microcentrifuge tubes. Each pellet was resuspended in 400 µl of lysis buffer and lysed using two rounds of sonication. The lysates were centrifuged (13,000 rpm, 10 min), and the supernatants were filtered through Amicon 10 kDa cut-off spin filters to remove proteins. A 96-well ELISA plate was washed five times with washing buffer. For the assay, 50 µl of each sample was transferred to the wells. Subsequently, 50 µl of c-di-AMP HRP tracer was added to each well except TA and blank wells, and 50 µl of c-di-AMP ELISA monoclonal antibody was added to each well except blank, TA, and NSB wells. The plate was incubated for two hours at room temperature with orbital shaking. After incubation, all wells were emptied and washed five times with wash buffer. Then, 175 µl of TMB substrate solution was added to each well and incubated for 30 minutes on an orbital shaker. Finally, 75 µl of HRP stop solution was added to each well, and absorbance was measured at 450 nm. The absorbance at 450 nm, measured spectrophotometrically, reflects the amount of cyclic di-AMP-HRP tracer bound to the well and is inversely proportional to the concentration of free c-di-AMP present during incubation.

### Plaque Assay

A recently developed protocol was followed^44^. Briefly, *H. volcanii* strains were grown overnight in Hv-ECa medium and normalized to an optical density of 595 nm (OD595). Next, 400 µl of cultures with 0.5 M CaCl2 was added to 3-4 ml of 0.2% top-agar (preheated and cooled to 60°C) in Hv-YPC, and the mixture was spread onto rich agar plates with 18% SW. The plates were incubated at room temperature for 15 to 20 minutes. A ten-fold serial dilution of HFPV-1 was prepared in Hv-ECa, and 3 µl of each dilution was spotted onto the agar. After 2 days of incubation at 30°C, plaques formed. The plates were then left on the bench at around 25°C for 2-3 days, during which time the plaques continued to grow and become more distinct. Images were captured to document the growth and appearance of the plaques.

### Long-term growth experiments

Individual colonies were selected from each strain (the *H. volcanii* strain containing H-TIR-SAVED and a control strain infected with the virus) and grown in liquid Hv-ECa medium supplemented with thymidine and tryptophan until reaching late logarithmic or stationary growth phase. The cultures were then subcultured into fresh medium, and this process was repeated for approximately 3 passages. After the third passage, 100 μl from each culture was serially diluted in fresh Hv-Ca medium and spread onto Hv-Ca plates containing thymidine and tryptophan. From the resulting growth, 25-30 colonies from each sample were analysed by PCR to assess HFPV-1 viral presence.

### Quantitative real-time PCR (Q-PCR)

We determined the genome copy numbers of *H. volcanii* and HFPV-1 using a CFX Connect Real-Time PCR system. Cultures were grown in triplicate for 72 hours, after which we collected 1 ml samples and centrifuged them at 11,000 × g for 10 minutes at room temperature. To measure viral DNA secretion, we extracted DNA from 200 μl of the resulting supernatant using a Quick-DNA Viral Kit. We then performed quantitative PCR with q-PCRBIO SyGreen Blue mastermix Hi-ROX, using primers targeting both an internal viral protein and the DNA polymerase II small subunit (*polB*), which served as a housekeeping gene. The viral DNA measurements were normalized by comparing their cycle threshold values to those of *polB*.

### Homology modelling using AlphaFold3

To create a homology model of the H-TIR-SAVED, we employed a multi-step process based on AlphaFold3 structure prediction. We first extracted the amino acid sequence of the H-TIR-SAVED (accession number: WP_115891642.1) from the NCBI database. The extracted sequence was submitted to the AlphaFold3 server for predicted structure generation. After obtaining the model, we conducted a Foldseek search for structural comparison against the PDB database. The resulting structure was analysed based on TM-score and RMSD (Root Mean Square Deviation) values. Finally, we used the UCSF Chimera software to visualize the alignments and generate the figures.

## Author contributions

U.G. and D.K.C. conceived and designed the study. D.K.C. performed the experiments with assistance from D.V. and H.S. D.K.C. also analyzed the data and prepared the figures. The manuscript was written by D.K.C. and U.G. All authors read and approved the final draft.

## Acknowledgements

The authors thank Prof. Sonja-Verena Albers for providing the strain with lower c-di-AMP levels and for discussions about c-di-AMP signalling in archaea. We thank Dr. Mantu Kumar for assistance with structure modeling and Alex Barbul for help with confocal microscopy. The authors also thank Dr. Leah Reshef for helpful discussions.

## References

1. Culbertson, E. M. & Levin, T. C. Eukaryotic CD-NTase, STING, and viperin proteins evolved via domain shuffling, horizontal transfer, and ancient inheritance from prokaryotes. PLOS Biol. 21, e3002436 (2023).

2. Millman, A., Melamed, S., Amitai, G. & Sorek, R. Diversity and classification of cyclic-oligonucleotide-based anti-phage signalling systems. Nat. Microbiol. 5, 1608–1615 (2020).

3. Tesson, F. et al. Systematic and quantitative view of the antiviral arsenal of prokaryotes. Nat. Commun. 13, 2561 (2022).

4. Cohen, D. et al. Cyclic GMP–AMP signalling protects bacteria against viral infection. Nature 574, 691–695 (2019).

5. Lowey, B. et al. CBASS Immunity Uses CARF-Related Effectors to Sense 3’-5’- and 2’-5’-Linked Cyclic Oligonucleotide Signals and Protect Bacteria from Phage Infection. Cell 182, 38–49.e17 (2020).

6. Duncan-Lowey, B., McNamara-Bordewick, N. K., Tal, N., Sorek, R. & Kranzusch, P. J. Effector-mediated membrane disruption controls cell death in CBASS antiphage defense. Mol. Cell 81, 5039–5051.e5 (2021).

7. Hogrel, G. et al. Cyclic nucleotide-induced helical structure activates a TIR immune effector. Nature 608, 808–812 (2022).

8. Rousset, F. et al. A conserved family of immune effectors cleaves cellular ATP upon viral infection. Cell 186, 3619–3631.e13 (2023).

9. Ledvina, H. E. et al. An E1–E2 fusion protein primes antiviral immune signalling in bacteria. Nature 616, 319–325 (2023).

10. Choudhary, D. K. et al. An archaeal CBASS system eliminates viruses without killing the host cells. 2024.09.12.612678 Preprint at 10.1101/2024.09.12.612678 (2024).

11. Govande, A. A., Duncan-Lowey, B., Eaglesham, J. B., Whiteley, A. T. & Kranzusch, P. J. Molecular basis of CD-NTase nucleotide selection in CBASS anti-phage defense. Cell Rep. 35, 109206 (2021).

12. Jacob, F. Evolution and Tinkering. Science 196, 1161–1166 (1977).

13. Molecular and Genome Evolution - Hardcover - Dan Graur - Oxford University Press. https://global.oup.com/ushe/product/molecular-and-genome-evolution-9781605354699?cc=il&lang=en.

14. Wang, S. et al. The role of TIR domain-containing proteins in bacterial defense against phages. Nat. Commun. 15, 7384 (2024).

15. Wani, K. A. & Pukkila-Worley, R. Evolutionarily ancient functions of enzymatic TIR proteins in innate immunity. Trends Immunol. 46, 441–454 (2025).

16. Burroughs, A. M., Zhang, D., Schäffer, D. E., Iyer, L. M. & Aravind, L. Comparative genomic analyses reveal a vast, novel network of nucleotide-centric systems in biological conflicts, immunity and signaling. Nucleic Acids Res. 43, 10633–10654 (2015).

17. Morehouse, B. R. et al. STING cyclic dinucleotide sensing originated in bacteria. Nature 586, 429–433 (2020).

18. Ofir, G. et al. Antiviral activity of bacterial TIR domains via immune signalling molecules. Nature 600, 116–120 (2021).

19. Shi, Y. et al. Structural characterization of macro domain–containing Thoeris antiphage defense systems. Sci. Adv. 10, eadn3310 (2024).

20. Ka, D., Oh, H., Park, E.Kim, J.-H. & Bae, E. Structural and functional evidence of bacterial antiphage protection by Thoeris defense system via NAD^+^ degradation. Nat. Commun. 11, 2816 (2020).

21. Slavik, K. M. & Kranzusch, P. J. CBASS to cGAS-STING: The Origins and Mechanisms of Nucleotide Second Messenger Immune Signaling. Annu. Rev. Virol. 10, 423–453 (2023).

22. Binder, S. C. et al. The SAVED domain of the type III CRISPR protease CalpL is a ring nuclease. Nucleic Acids Res. 52, 10520–10532 (2024).

23. Steens, J. A., Salazar, C. R. P. & Staals, R. H. J. The diverse arsenal of type III CRISPR-Cas-associated CARF and SAVED effectors. Biochem. Soc. Trans. 50, 1353–1364 (2022).

24. Rouillon, C. et al. Antiviral signalling by a cyclic nucleotide activated CRISPR protease. Nature 614, 168–174 (2023).

25. Athukoralage, J. S. & White, M. F. Cyclic Nucleotide Signaling in Phage Defense and Counter-Defense. Annu. Rev. Virol. 9, 451–468 (2022).

26. Alarcón-Schumacher, T., Naor, A., Gophna, U. & Erdmann, S. Isolation of a virus causing a chronic infection in the archaeal model organism Haloferax volcanii reveals antiviral activities of a provirus. Proc. Natl. Acad. Sci. U. S. A. 119, e2205037119 (2022).

27. Giani, M., Miralles-Robledillo, J. M., Peiró, G., Pire, C. & Martínez-Espinosa, R. M. Deciphering Pathways for Carotenogenesis in Haloarchaea. Mol. Basel Switz. 25, 1197 (2020).

28. Kellermann, M. Y., Yoshinaga, M. Y., Valentine, R. C., Wörmer, L. & Valentine, D. L. Important roles for membrane lipids in haloarchaeal bioenergetics. Biochim. Biophys. Acta BBA - Biomembr. 1858, 2940–2956 (2016).

29. Vögeli, B. et al. Archaeal acetoacetyl-CoA thiolase/HMG-CoA synthase complex channels the intermediate via a fused CoA-binding site. Proc. Natl. Acad. Sci. 115, 3380–3385 (2018).

30. Abramson, J. et al. Accurate structure prediction of biomolecular interactions with AlphaFold 3. Nature 630, 493–500 (2024).

31. Abramson, J. et al. Accurate structure prediction of biomolecular interactions with AlphaFold 3. Nature 630, 493–500 (2024).

32. Ofir, G. et al. Antiviral activity of bacterial TIR domains via immune signalling molecules. Nature 600, 116–120 (2021).

33. Braun, F. et al. Cyclic nucleotides in archaea: Cyclic di-AMP in the archaeon Haloferax volcanii and its putative role. MicrobiologyOpen 8, e00829 (2019).

34. van der Does, C., Braun, F., Ren, H. & Albers, S.-V. Putative nucleotide-based second messengers in archaea. microLife 4, uqad027 (2023).

35. Galperin, M. Y. All DACs in a Row: Domain Architectures of Bacterial and Archaeal Diadenylate Cyclases. J. Bacteriol. 205, e00023–23 (2023).

36. DeWeirdt, P. C., Mahoney, E. M. & Laub, M. T. DefensePredictor: A Machine Learning Model to Discover Novel Prokaryotic Immune Systems. 2025.01.08.631726 Preprint at 10.1101/2025.01.08.631726 (2025).

37. Hobbs, S. J. et al. Phage anti-CBASS and anti-Pycsar nucleases subvert bacterial immunity. Nature 605, 522–526 (2022).

38. Cao, X. et al. Phage anti-CBASS protein simultaneously sequesters cyclic trinucleotides and dinucleotides. BioRxiv Prepr. Serv. Biol. 2023.06.01.543220 (2023) doi:10.1101/2023.06.01.543220.

39. Makarova, K. S. et al. Evolutionary and functional classification of the CARF domain superfamily, key sensors in prokaryotic antivirus defense. Nucleic Acids Res. 48, 8828–8847 (2020).

40. Whiteley, A. T. et al. c-di-AMP modulates Listeria monocytogenes central metabolism to regulate growth, antibiotic resistance and osmoregulation. Mol. Microbiol. 104, 212–233 (2017).

41. Oppenheimer-Shaanan, Y., Wexselblatt, E., Katzhendler, J., Yavin, E. & Ben-Yehuda, S. c-di-AMP reports DNA integrity during sporulation in Bacillus subtilis. EMBO Rep. 12, 594–601 (2011).

42. Zarrella, T. M. & Bai, G. The Many Roles of the Bacterial Second Messenger Cyclic di-AMP in Adapting to Stress Cues. J. Bacteriol. 203, e00348–20 (2020).

43. Govande, A. A., Duncan-Lowey, B., Eaglesham, J. B., Whiteley, A. T. & Kranzusch, P. J. Molecular basis of CD-NTase nucleotide selection in CBASS anti-phage defense. Cell Rep. 35, 109206 (2021).

44. Navok, S., Cohen, L., Ron, E. Z. & Gophna, U. Optimized Plaque Assay for Detecting Chronically Infecting Viruses of Haloarchaea. 2024.11.12.619373 Preprint at 10.1101/2024.11.12.619373 (2024).

