## Extended figures for "A TIR-SAVED effector mediates antiviral immunity via a conserved host signal"

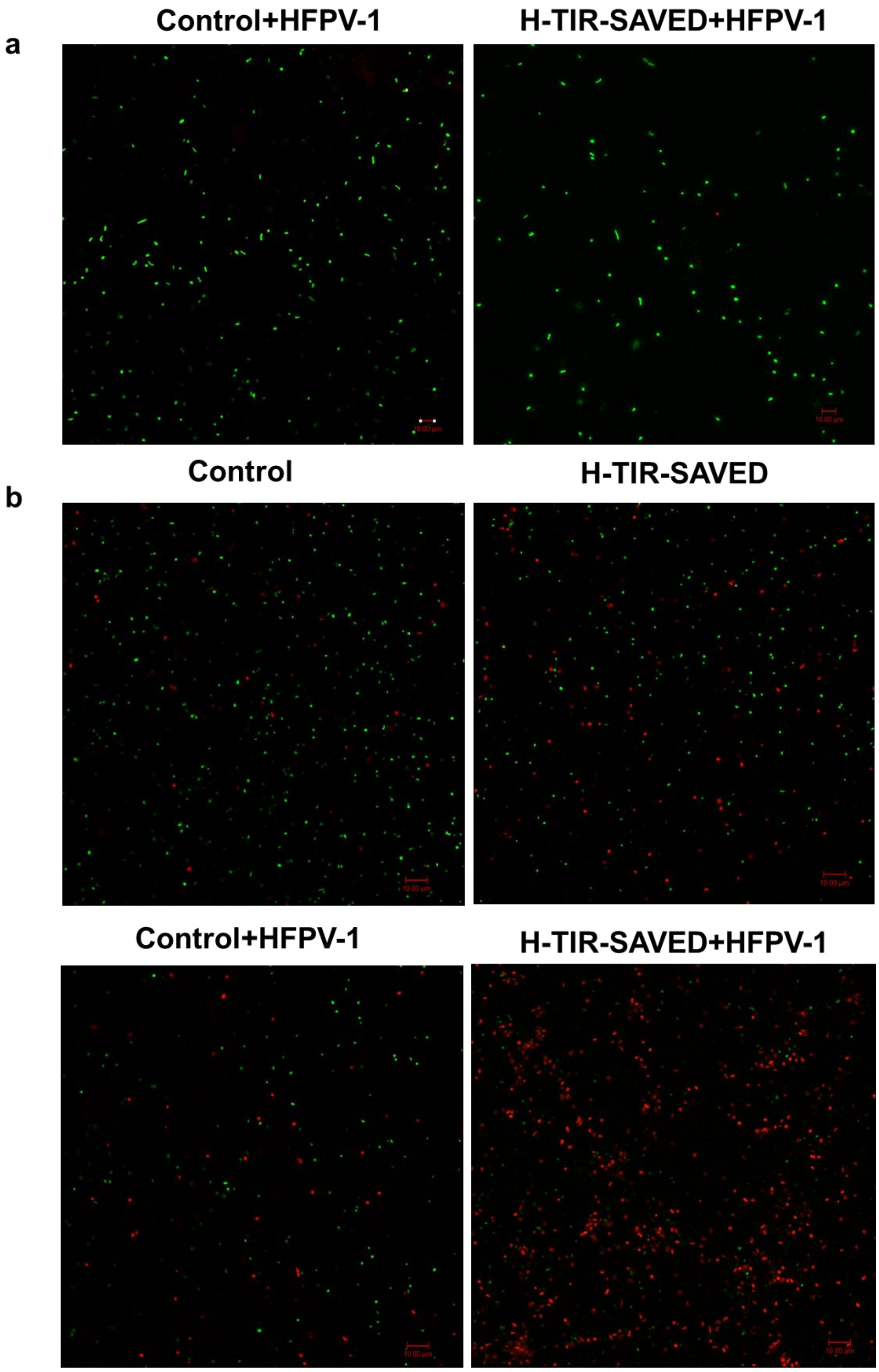


**Extended Fig. 1. H-TIR-SAVED does not induce rapid cell death following HFPV-1 infection.** a) Representative confocal microscopy images from Live/Dead assays of log-phase *H. volcanii* cells. Cells expressing H-TIR-SAVED and control cells carrying an empty vector were infected with HFPV-1. No significant cell death was observed in virus-infected log-phase cells expressing H-TIR-SAVED (Scale bar: 10 µm). Data represent at least three independent experiments. b) Live/Dead assays of cells from bleached colonies (maintained at room temperature for 4–5 weeks) showed an increased number of dead cells in the virus-infected strain expressing H-TIR-SAVED compared to the control (Scale bar: 10 μm).

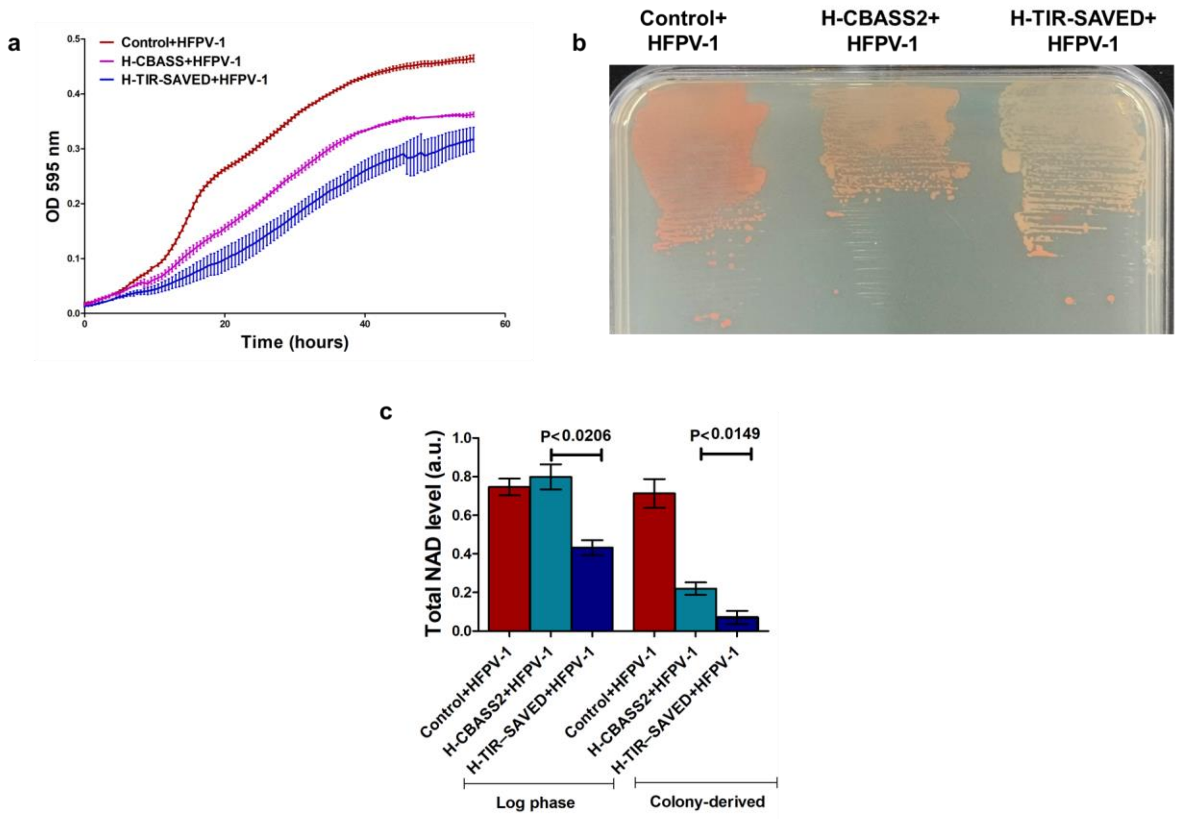


**Extended Fig. 2 Virus infection induces more severe growth arrest in H-TIR-SAVED–expressing cells than in cells expressing the full-length H-CBASS2 system**. a) Growth curves of virus-infected *H. volcanii* strains expressing H-CBASS2 (purple), H-TIR-SAVED (blue), and the infected control (red). Values represent the mean ± SE of at least three biological replicates, each with three technical replicates. b) Bleaching of virus-infected H-TIR-SAVED and H-CBASS2-expressing *H. volcanii* colonies. Representative images show colony color of HFPV-1–infected *H. volcanii* following plate streaking (see Methods). H-TIR-SAVED-expressing colonies displayed more rapid bleaching after 3–4 weeks at room temperature compared to H-CBASS2-expressing colonies. c) $NAD^+$ levels in *H. volcanii* strains expressing H-TIR-SAVED and H-CBASS2 were measured following virus infection, from both log phase cultures and bleached colonies. $NAD^+$ levels depleted more rapidly in H-TIR-SAVED-expressing cells compared to full-length H-CBASS2-expressing cells upon virus infection. Data represent results from three independent experiments.

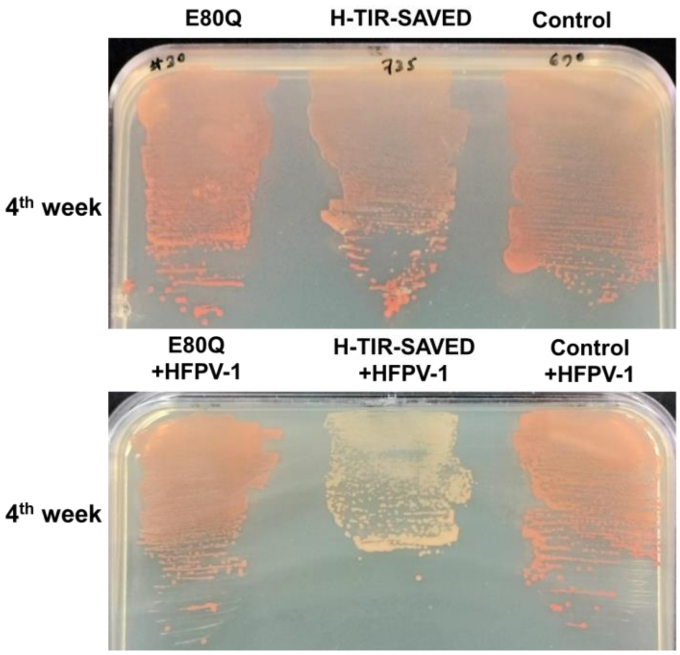


**Extended Fig. 3) The E80Q mutation in TIR-SAVED did not result in bleaching after prolonged incubation.** A plate streaking assay was conducted using *H. volcanii* strains expressing E80Q/H-TIR-SAVED, and TIR-SAVED both with and without viral infection, along with a control containing an empty vector. Images were captured after 3-4 weeks of incubation as described previously.

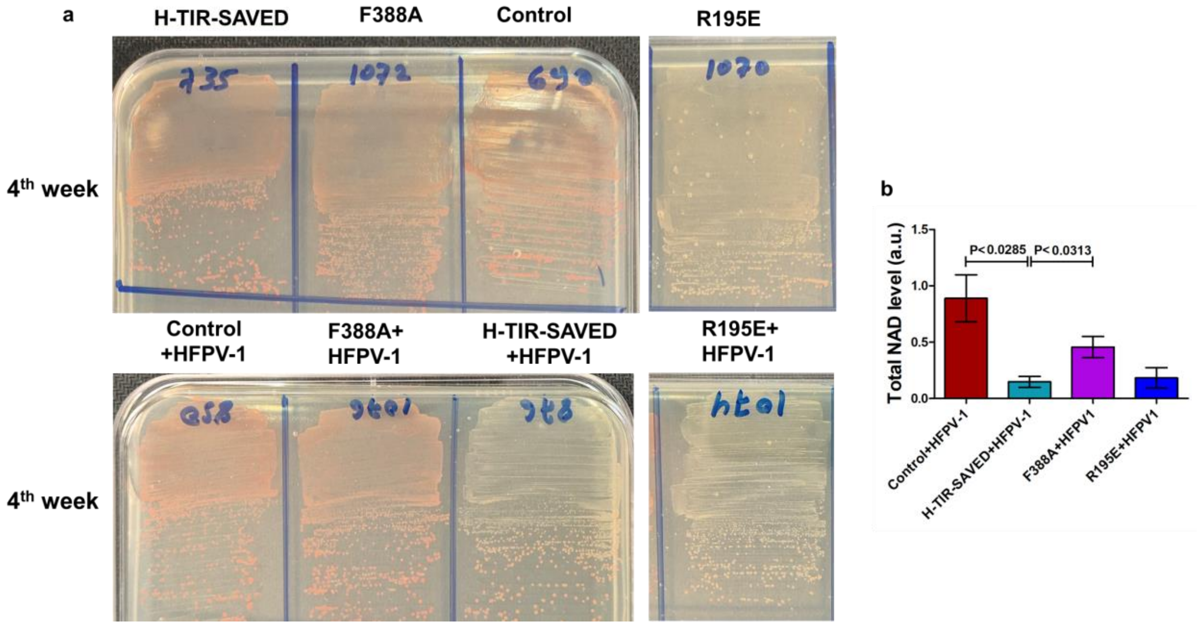


**Extended Fig. 4) The F388A mutation in TIR-SAVED resulted in very slow bleaching after prolonged incubation.** a) A plate streaking assay was performed using *H. volcanii* strains expressing F388A/H-TIR-SAVED, R195E/H-TIR-SAVED, and TIR-SAVED, both with and without viral infection, along with a control containing an empty vector. Images were captured after 3-4 weeks of incubation as described previously. b) $NAD^+$ levels in *H. volcanii* strains expressing F388A and R195E mutations in the TIR-SAVED protein. Strains were grown to stationary phase, and $NAD^+$ levels were measured following the described protocol. Data are representative of three independent experiments. Statistical analysis was performed using a paired t-test.

**Extended table 1. List of all the strains and plasmids used in this study**

| Archaeal strains & plasmids | Description | Source / Reference |
|---|---|---|
| WR540 | *H.volcanii ΔpyrE2, ΔhdrB, ΔtrpA* | (Allers et al., 2004) |
| WR532 | *H.volcanii* ΔpyrE2 | (Allers et al., 2004) |
| UG687 | WR540 + CBASS type II cloned in pTA927 | (Choudhary et al., 2024) |
| UG690 | WR540 + pTA927 | (Choudhary et al., 2024) |
| UG850 | UG690+ HFPV-1 | (Choudhary et al., 2024) |
| UG851 | UG687+ HFPV-1 | (Choudhary et al., 2024) |
| UG840 | WR532 + HFPV-1 | (Choudhary et al., 2024) |
| UG735 | WR540 + H-TIR-SAVED cloned in pTA927 | This study |
| UG876 | UG735+HFPV-1 | This study |
| UG906 | WR540+E80Q mutation on H-TIR-SAVED cloned in pTA927 | This study |
| UG1136 | UG906+HFPV-1 | This study |
| UG908 | WR540 + H-E1E2-JAB cloned in pTA927 | This study |
| UG1134 | UG908+HFPV-1 | This study |
| UG1070 | WR540 +R195E mutation on H-TIR-SAVED cloned in pTA927 | This study |
| UG1074 | UG1070+HFPV-1 | This study |
| UG1072 | WR540 +F388A mutation on H-TIR-SAVED cloned in pTA927 | This study |
| UG1076 | UG1072+HFPV-1 | This study |
| UG992 | HTQ410 (*H98, ΔpyrE2, ΔhdrB, p.tnaM3-dacZ-hdrB+*) | Braun et al.,2019 |
| UG1017 | UG992+ pTA927 | This study |
| UG1035 | UG1017+ HFPV-1 | This study |

| | | |
|---|---|---|
| UG1023 | UG992+ H-TIR-SAVED cloned in pTA927 | This study |
| UG1039 | UG1023+HFPV-1 | This study |
| UG1216 | UG992+ H-CBASS2 cloned in pTA927 | This study |
| UG1240 | UG1216+HFPV-1 | This study |
| **Plasmid** | **Description** | **Source / Reference** |
| pTA927 | Expression vector with *pyrE2* marker and pHV2 origin. | [33] |
| pUG683 | H-CBASS2 cloned into pTA927/*E. coli* | (Choudhary et al., 2024) |
| pUG745 | H-TIR-SAVED cloned into pTA927/ *E. coli* | This study |
| pUG934 | E80Q mutation on H-TIR-SAVED cloned in pTA927/ *E. coli* | This study |
| pUG1083 | R195E mutation on H-TIR-SAVED cloned in pTA927/ *E. coli* | This study |
| pUG1086 | F388A mutation on H-TIR-SAVED cloned in pTA927/ *E. coli* | This study |

**Extended table 2. List of oligonucleotides used in this study**

| Primer | Sequence (5’→ 3’) | Description |
|---|---|---|
| DKC49 | GCGGACCTATTGCGCATATGACTAACCCCACTGGAGAGGT | Forward primers for H-TIR-SAVED (insert) cloning in pTA927 |
| DKC50 | GATGGTCCAGAGGTGCGGCCCTATTGTATCAGGATGGCTG | Reverse primers for H-TIR-SAVED (insert) cloning in pTA927 |
| DKC51 | CAGCCATCCTGATACAATAGGGCCGCACCTCTGGACCATC | Forward primers for H-TIR-SAVED (plasmid) cloning in pTA927 |
| DKC52 | ACCTCTCCAGTGGGGTTAGTCATATGCGCAATAGGTCCGC | Reverse primers for H-TIR-SAVED (plasmid) cloning in pTA927 |

| | | |
|---|---|---|
| DKC127 | CCTAAACGTAcagATCCCCCAAATAC | Forward primer for the mutation on TIR-SAVED domain at E80Q |
| DKC128 | ATTGTTGGCGAATCGAGG | Reverse primer for the mutation on TIR-SAVED domain at E80Q |
| DKC141 | CGAAGACAATgaaCTTCCAGGAGAAAAAAC | Forward primer for the mutation on TIR-SAVED domain at R195E |
| DKC142 | AAGTGGTTGGATAGGTCG | Reverse primer for the mutation on TIR-SAVED domain at R195E |
| DKC143 | AACATACGCGgctAATGACAACCATCAG | Forward primer for the mutation on TIR-SAVED domain at F388A |
| DKC144 | TGGATTGGAGGTAGCGTG | Reverse primer for the mutation on TIR-SAVED domain at F388A |
| DKC95 | TTGCGTACGCGGTATCTGTC | Forward primer for the q-PCR of HFPV-1 |
| DKC96 | AGCTTCTCCGCATCGTCTTT | Reverse primer for the q-PCR of HFPV-1 |
| DKC97 | CACGAACGAGAACACCGACC | Forward primer for the detection of virus (HFPV-1) |
| DKC98 | TGATGACGAATCCAACGAGCAG | Reverse primer for the detection of virus (HFPV-1) |
| DKC157 | CCCGAATCAGGACGAAGAAC | Forward primer for the q-PCR of polB |
| DKC158 | ATTTGAGGTGCTCGGAGAAC | Reverse primer for the q-PCR of polB |
